# Arthropod succession from deadwood to soil: increasing diversity and transitions between major forest substrates

**DOI:** 10.64898/2026.09.21.753076

**Authors:** William Esbjug Gromstad, Tone Birkemoe, Håvard Kauserud, Johan Asplund, Sundy Maurice, Eivind Kverme Ronold, Celina Nilsen, Anders K. Krabberød, Lisa Fagerli Lunde

## Abstract

Deadwood is a common ephemeral resource in boreal forests, but arthropod succession across the full decay gradient and transition into soil remains poorly understood. We analyzed the successional trajectory of arthropods in deadwood to soil. Further, we investigated whether the succession pattern differed in previously clear-cut and near-natural forests in southeastern Norway. This was done by DNA metabarcoding analyses of arthropod communities in collected deadwood and soil samples. We detected 741 arthropod OTUs, with 565 OTUs in deadwood and 291 in organic soil layers, of which 115 were found in both deadwood and organic soil layers. Arthropod richness increased with deadwood decay stage and organic soil layers supported higher richness than all deadwood decay stages. Community composition of arthropods changed significantly along the decay gradient, with late-decay stages sharing more OTUs with organic soil layers than with earlier decay stages. This suggests a gradual transition between communities in highly decayed deadwood and organic soil layers. Forest type (previously clear-cut vs. near-natural forest) had limited effects on overall richness and community composition, although previously clear-cut and near-natural forests supported partially distinct species pools and differed in co-occurrence patterns in late decay and organic soil layers. Our results show that arthropod communities are strongly structured along the decay gradient from freshly dead wood to organic soil layers. This highlights the importance of retaining deadwood across different decay stages for maintaining arthropod diversity and community convergence between deadwood and soil in boreal forests.

## Introduction

Ecological succession is a fundamental process in which biological communities change over time in response to environmental conditions and resource availability (Connell and Slatyer, 1977, Fukami, 2015, Chang and Turner, 2019). Successional dynamics occur across a wide range of temporal and spatial scales, from forest succession following large-scale disturbances such as wildfire to succession within ephemeral resource patches that develop and disappear within a relatively short time (Connell and Slatyer, 1977, Walker and del Moral, 2003, Chang and Turner, 2019). According to successional theory, temporal changes in habitat conditions modify resource availability and species interactions (Connell and Slatyer, 1977, Fukami, 2015). As a result, species colonization and community assembly depend on the successional stage of the resource (Connell and Slatyer, 1977, Walker and del Moral, 2003, Fukami, 2015).

Deadwood represents a common ephemeral resource in forests, undergoing major physical and chemical transformations during decay that drive shifts in associated biological communities (Stokland et al., 2012, Vindstad et al., 2020). At the same time, deadwood supports a substantial proportion of forest biodiversity, as many arthropods, fungi, and other microorganisms depend upon deadwood for at least one part of their lifecycle (Lassauce et al., 2011, Stokland et al., 2012, Seibold et al., 2015). Habitat heterogeneity within deadwood is expected to increase during decay, as physical and chemical changes create an increasingly diverse range of microhabitats and resources. This expectation is supported by previous research showing that arthropod and fungal richness increase with advancing decay (Kim et al., 2013, Mäkipää et al., 2017). When decay progresses, deadwood becomes increasingly integrated into the forest floor, as gradual physical breakdown and nutrient release create a continually changing habitat, ultimately with similar properties as soil (Ulyshen, 2016, Mäkipää et al., 2017). Consequently, late-decay deadwood is expected to support an increasing proportion of soil-associated taxa, as habitat conditions over time become more similar to those in the organic soil layer, leading to higher overlap between communities. Understanding succession in deadwood can therefore reveal how the continuous development of a habitat influences species accumulation and community composition over time. Arthropods are particularly well-suited for studying these transitions because they occupy a wide range of trophic roles and are closely linked to changes in habitat structure and resource availability (Mäkipää et al., 2017). While beetles in deadwood are relatively well researched, most other deadwood-associated arthropods are not (Stokland et al., 2012, Parisi et al., 2018).

The succession of arthropods follows a relatively well-defined trajectory (Stokland et al., 2012, Parisi et al., 2018). In boreal forests, Norway spruce (*Picea abies*) deadwood succession has been particularly well studied (Stokland et al., 2012). Five broad successional stages are commonly recognized (Renvall, 1995). In the first decay stage (DW1), recently dead trees are primarily colonized by specialists such as bark beetles and other phloem-feeding insects such as weevils and longhorn beetles, with their associated predators (Stokland et al., 2012). These early colonizers may initiate deadwood decay directly by initiating bark loss (Renvall, 1995), and indirectly by bringing decay fungi into the wood (Jacobsen et al., 2017, Lunde et al., 2023). In the second stage (DW2), bark begins to loosen and fungal mycelia is well established. Thus, fungivorous arthropods often dominate the subcortical habitat between the bark and the wood (Stokland et al., 2012). The third stage (DW3) begins as the subcortical habitat disappears when the bark detaches and fungi become increasingly abundant (Boddy, 2001, Baldrian, 2008). Decay fungi drive the enzymatic breakdown of cellulose, hemicellulose, and lignin (Stokland et al., 2012), thereby facilitating colonization by fungivorous and detritivorous arthropods (Stokland et al., 2012, Parisi et al., 2018). In the final decay stages (DW4 and 5), the wood is largely decomposed, and the habitat becomes increasingly similar to soil, which is reflected by the presence of more generalist detritivores and soil-associated arthropods (Stokland et al., 2012, Mäkipää et al., 2017). Because the complete decay of deadwood is slow in boreal forests, these later decay stages enable the development of long-term interactions between deadwood and soil communities (Harmon et al., 1986). In addition to the species typical for specific decay stages, their associated predators are present in all stages (Stokland et al., 2012).

Forest management may impact arthropod succession in deadwood by altering the composition of deadwood in a forest stand and thus shaping the local species pools (Parisi et al., 2018, Lunde et al., 2025). For instance, previously clear-cut boreal forests often have lower volumes and diversity of deadwood in comparison to near-natural forests (Asplund et al., 2024, Asplund et al., 2026), which directly reduces habitat availability for deadwood-associated arthropods. Because these forests often lack large, late-decay deadwood (Asplund et al., 2024, Lunde et al., 2025), the gradual ecological transition between deadwood and soil, and their associated communities, may also be disrupted. Thus, understanding how forestry practices alter successional dynamics is important for predicting arthropod diversity, community composition, and ecological transitions between deadwood and soil communities across forest substrates.

Despite the ecological importance of deadwood-associated communities, our understanding of arthropod successional dynamics remains incomplete. Traditional approaches have provided valuable insights into the deadwood-associated fauna using emergence traps, eclector traps, wood dissection, and extensive morphological identification (Stokland et al., 2012, Seibold et al., 2015, Ulyshen, 2016). However, these approaches are often time-consuming and focus on specific taxa such as beetles (Stokland et al., 2012, Seibold et al., 2015, Ulyshen, 2016). As a result, relatively few studies have examined the succession of the entire arthropod community across the full deadwood decay gradient (but see Kim et al., 2013). DNA metabarcoding has emerged as a powerful tool for overcoming these limitations, enabling the detection of broader and cryptic taxa (Taberlet et al., 2012, Deiner et al., 2017), although so far with variable success for deadwood-associated arthropods (Winiger et al., 2022).

In this study, we used DNA metabarcoding to investigate arthropod successional trajectories across the deadwood decay gradient and their convergence with organic soil. We also examined whether these successional trajectories differed between previously clear-cut and near-natural boreal forests. Specifically, we tested the following hypotheses: H1. Arthropod diversity increases along the deadwood decay gradient. H2. Arthropod community composition changes predictably along the decay gradient, with late-decay communities becoming increasingly similar to those in organic soil layers. H3. Previously clear-cut and near-natural forests differ in arthropod diversity, community composition, and the overlap between communities in deadwood and organic soil layers.

## Materials and methods

### Site description and study design

Our study was conducted in boreal forests in southeastern Norway, across 12 site pairs ranging 200 km in latitude and 160 km in longitude (Figure 1). The sites were dominated by *Picea abies* (Norway spruce), with understories primarily consisting of bilberry (*Vaccinium myrtillus*) and feather mosses such as *Hylocomium splendens* and *Pleurozium schreberi*. Mean annual temperatures varied between 0.5 to 5.4 °C, and mean annual precipitation ranged from 683 to 1056 mm. Each site pair consisted of two forest types, one previously clear-cut forest and one near-natural forest, as identified in the EcoForest project and described in detail in Asplund et al., 2024. Previously clear-cut forests belonged to maturity classes IV–V (Breidenbach et al., 2020), with an average breast height-age of dominating trees between 43 and 82 years. The near-natural forests were older, with trees averaging between 99 and 189 years old. These forests have previously only been exposed to selective logging, but had few characteristics related to recent human activities (Asplund et al., 2024).

**Figure 1.**
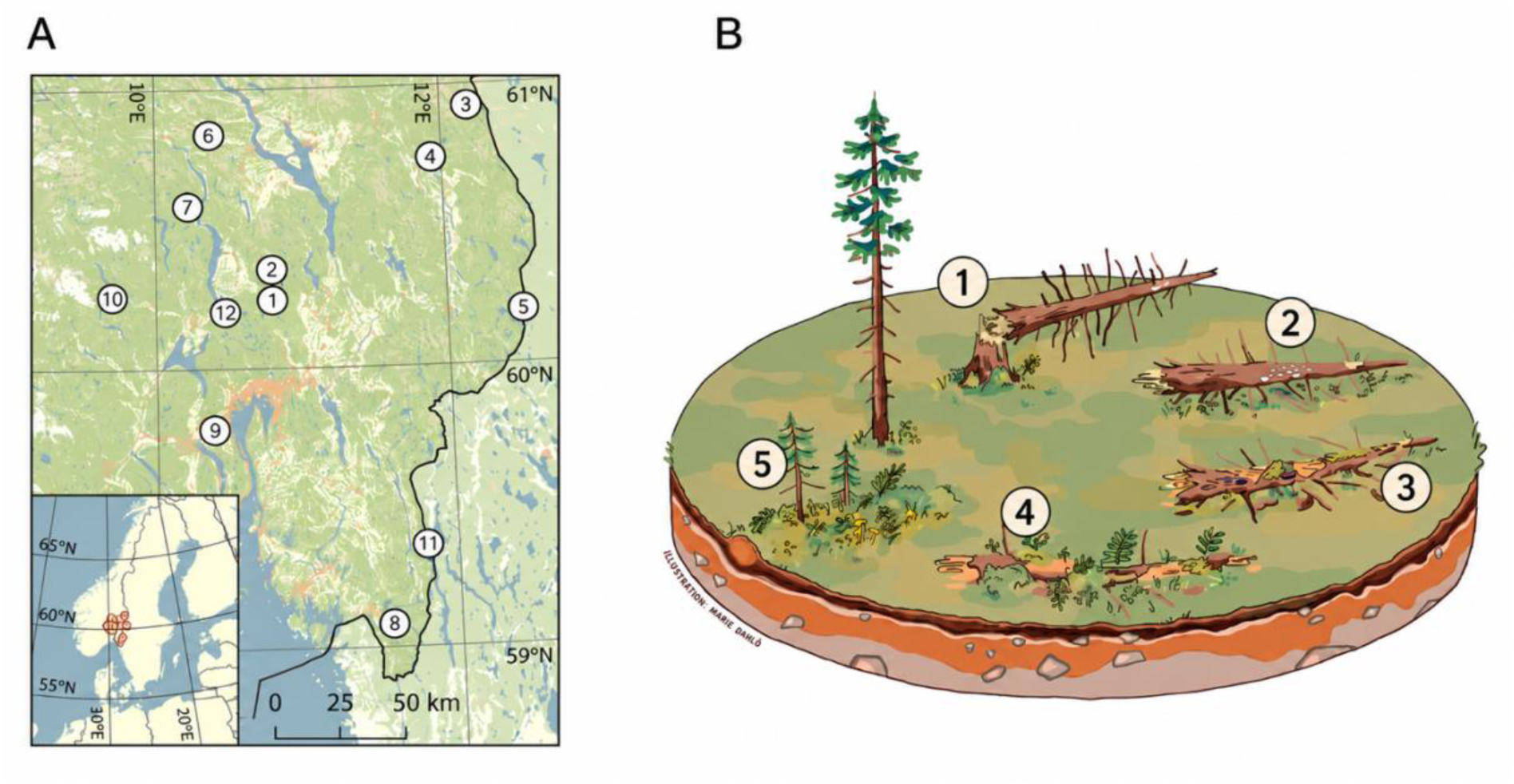
(A) Location of the twelve study sites in southeastern Norway. Each site consisted of one previously clear-cut forest and one near-natural forest. Reproduced from Asplund et al. (2024) under CC BY 4.0. (B) Visualization of deadwood decay stages: (1) Recently dead wood with bark firmly attacked or recently loosened after bark beetle attack, but wood still hard. (2) Bark loose with developing fungal mycelium. The wood can be penetrated up to 3 cm with a knife. (3) Advanced decay with mostly soft wood, but the inner core remains hard. (4) Wood decomposed throughout and can be broken apart by hand. (5) Highly decomposed wood fragments, with only remnants of the original log structure remaining. Illustrated by Marie David.

To identify arthropods in deadwood, we established a 15 × 133 m transect within each site. Twenty deadwood logs (over 5 cm in diameter at the base) of Norway spruce were randomly chosen within each transect for sawdust collection. Some sites had less than 20 logs of deadwood. In these cases, all available deadwood logs were chosen, leading to a sawdust collection from 471 deadwood logs assigned to decay stages on a 0–5 scale (Table 1). There were differences in total log volume between forest types, and the near-natural forests had larger logs, especially for later decay stages (Asplund et al., 2024).

**Table 1:** Number of deadwood logs within different decay stages in total, in previously clear-cut (CC) and near-natural (NN) forests, and the mean log volume and diameter with standard deviation of logs in both forest types.

| Decay stage | Logs | Logs in CC/NN | CC mean volume (m<sup>3</sup>)<br>± SD | NN mean volume (m<sup>3</sup>)<br>± SD | CC mean diameter (cm) ± SD | NN mean diameter (cm) ± SD |
| --- | --- | --- | --- | --- | --- | --- |
| 1 | 142 | 81/61 | 0.015 ± 0.028 | 0.040 ± 0.142 | 7.93 ± 2.71 | 9.18 ± 5.03 |
| 2 | 158 | 61/97 | 0.040 ± 0.066 | 0.153 ± 0.265 | 11.10 ± 5.07 | 14.38 ± 8.63 |
| 3 | 76 | 40/36 | 0.032 ± 0.059 | 0.207 ± 0.278 | 9.42 ± 4.31 | 17.50 ± 10.81 |
| 4 | 50 | 28/22 | 0.026 ± 0.023 | 0.237 ± 0.287 | 12.10 ± 5.47 | 21.09 ± 12.49 |
| 5 | 17 | 9/8 | 0.062 ± 0.114 | 0.116 ± 0.117 | 16.22 ± 8.07 | 20.22 ± 9.07 |

Sawdust samples were collected to target arthropod DNA in deadwood. Before drilling, the outer bark and the 1–2 millimeter of the surface wood were removed in order to only collect sawdust from inside the log. Sawdust was then collected by drilling horizontally 6 cm deep with a 22 mm drill bit. For small logs (< 6 cm diameter), drilling was stopped before penetrating the opposite side of the log. To better cover spatial heterogeneity within logs, samples were collected either at the center of short logs (< 2.5 m) or at two m intervals along longer logs, starting 50 cm from the root collar or point of break when not uprooted. Samples were stored on ice during field days (1–3 days), and subsequently frozen at −20°C until DNA extraction.

To study the potential convergence between deadwood and soil arthropods, soil samples (ø = 27 mm and 30 cm depth) were taken in a 25-point systematic grid. When rocks or roots hindered exact sampling positions, soil cores were collected as close as possible to the intended grid-point. Each soil core was then divided into seven layers based on functionality: litter/fermentation (LF), fermentation/humus (FH), A layer (divided into the top five cm and the rest), AB transition, B layer (divided into the top five cm and the rest). Each soil layer was pooled into one sample per site, resulting in one composite sample for each layer from each plot. Approximately 2 g of homogenized, freeze-dried, and crushed soil from each pooled sample was used for DNA analysis. The seven soil layers were separated into organic (LF and FH) and mineral (rest) soil after DNA metabarcoding analysis. However, only organic soil layers were used in further data analysis due to low sequencing depth in mineral soil layers.

### Environmental variables

To characterize environmental conditions potentially influencing arthropod succession, several abiotic and structural variables were measured for both deadwood and soil samples. Variables related to moisture, nutrient status, acidity, and substrate size were included as covariates in subsequent analyses.

Deadwood volume (m^3^) was estimated from log length and diameter measurements and included as a proxy for habitat size and substrate availability (Table 1). Moisture content (%) was measured from wood samples collected during field sampling. In addition, pH and carbon-to-nitrogen (C:N) ratio were included to represent chemical changes occurring during decay and their potential influence on fungal and arthropod communities. For soil samples, pH, soil bulk density (g L^-1^), and C:N ratio were included as environmental variables because these are known to influence soil arthropod communities and microbial activity. Mean annual temperature (MAT) and slope were additionally included to account for climatic and topographic variation among sites.

### Sample processing, DNA extraction and metabarcoding

Deadwood and soil samples were sampled separately and processed differently, as described in Ronold et al., 2026a and 2026b. Deadwood samples were homogenized by thoroughly mixing the sawdust inside each plastic bag. Around ∼6 g of fresh sawdust was transferred into a pre-weighted 50 ml Falcon Tube. The sawdust was freeze-dried for 48 h until there was no more weight reduction. Moisture content was determined from the difference in sample mass before and after freeze-drying. The dried sawdust was equally divided into two new Falcon Tubes for DNA extractions and chemical analyses. Sawdust samples were pulverized by adding two 5 mm ceramic beads to each tube and processing the samples in a FastPrep 24 instrument (MP Biomedicals, USA) at 4.5 m s⁻¹ for 45 s. This was repeated a minimum of three times per tube. Sawdust from early decay logs required repeated runs to fully break apart the wood chips up to nine times per tube at the same speed. DNA was extracted using an optimized in-house CTAB-chloroform protocol for a relatively large amount of sawdust, followed by an additional DNA-column cleaning step. The DNA extracts were cleaned using an E.Z.N.A 96-well plate kit following the manufacturer’s protocol (Omega Bio-tek) and the elute were collected in 96-well PCR microplates stored at –20°C.

DNA from soil was extracted using an E.Z.N.A 96-well plate kit (Omega BioTek) following the manufacturer’s protocol (Ronold et al., 2026b). Briefly 600 µl CTAB buffer was added to dried, crushed soil and vortexed. A 725 µl sample of homogenized slurry was transferred to an E-Z 96-well Disruptor Plate C Plus, sealed and processed on a Tissue Lyser II (Qiagen) for two cycles of 2.30 min at 20 rpm, with inversion between cycles. DS Buffer (72 µl) was added, and plates were vortexed and incubated at 70 °C for 10 min with intermittent mixing. After centrifugation at 2000g for 10 mins, 300 µl of supernatant was transferred to a new 96 deep-well plate. DNA was purified using the E.Z.N.A 96 well kit according to the manufacturer’s protocol, eluted in 100 µl and stored at −20°C.

From both deadwood and soil, we amplified a ∼ 421 bp fragment of the COI gene using the BF3/BR2 primer pair, designed to broadly target arthropod DNA while maintaining high taxonomic resolution (Elbrecht et al., 2019). To enable multiplexed sequencing and reduce the risk of incorrect assignment of samples, each primer contained a unique seven-nucleotide tag at the 5’ end. The PCR mix contained 5 ml Q5 buffer, 5 ml Q5 enhancer, 1 ml (20 mg ml^-1^) BSA, 0.5 ml dNTP (10 mM), 9.75 ml H_2_O, 1.2 ml BF3 primer, 1.2 ml BR2 primer, 1 ml DNA template and 0.25 ml polymerase. The PCR run for DNA amplification included an initial phase of denaturation and enzyme-activation for 5 minutes at 98°C, followed by 30 cycles of 98°C for 30 seconds, an annealing phase at 50°C for 30 seconds and extension phase at 72°C for 50 seconds, before a final extension phase at 72°C for 10 minutes. In addition to the COI amplicons, we had one biological replicate, one mock sample, one PCR negative, two extraction negatives, four internal plate replicates, and seven cross-plate replicates in each plate. Including both soil and deadwood samples, eight plates underwent the PCR process. PCR amplicons were normalized using SequalPrep Normalization Plate Kit, 96-well (Thermo Fisher Scientific) following the manufacture’s protocol. Amplicons were further cleaned with AMPure XP Beads (Backman Coutler). For each 96-well SequalPrep Normalization plate, 4 columns of PCR product were pooled in 1.5 mL Eppendorf tube, resulting in 3 pools of product. The volume of each tube was measured and 0.8X volume of AMPure Beads was added. Tubes were mixed on Eppendorf Thermomixer comfort at maximum speed for 10 minutes and then placed on magnetic rack for 5 minutes. Supernatant was removed, twice 500 µl of 80% ethanol was added and left for 2 minutes before being removed, although on the second removal only 470 µl was removed. The beads were then spun down, replaced on magnetic rack to remove the remaining ethanol before adding 20 µl EB buffer. The 3 tubes were then pooled together into one final ∼60 µl library in a clean 1.5 mL Eppendorf tube. In total 9 libraries were normalized and cleaned. DNA concentrations were measured using Qubit BR DNA Assay (Thermo Fisher Scientific). All libraries were sent to be sequenced on the Illumina MiSeq platform at Fasteris (Plan-Les-Ouates (GE), Switzerland).

### Bioinformatics

Bioinformatic processing of the COI data was performed on the SAGA computer cluster provided by Sigma2, the National Infrastructure for High-Performance Computing and Data Storage in Norway, using a custom pipeline (https://github.com/ekronold/Zazzy_metabarcoding_pipeline). Paired-end reads were demultiplexed and primers were removed using Cutadapt v4.2 (Martin, 2011), with a minimum sequence length of 100 bp and a required 26-bp overlap between the tags and primer sequences. One mismatch was allowed in the tags and only sequences with matching forward and reverse tags were retained. Sample files containing fewer than 5000 characters were discarded. Demultiplexed samples were quality-filtered, denoised, and checked for chimeras using DADA2 (Callahan et al., 2015). In filterAndTrim(), maxEE was set to c(4, 4), and reads were truncated using truncLen = c(265, 215). During denoising, the global option BAND_SIZE was set to 16. Chimera removal was performed using the default settings. The resulting ASVs were clustered into OTUs at 97% sequence similarity using VSEARCH v2.22 (Rognes et al., 2016). A second de novo chimera check was performed with VSEARCH to remove chimeras potentially missed by DADA2 (--uchime_denovo). The OTU table was subsequently curated using the mumu post-clustering algorithm (https://github.com/frederic-mahe/mumu) to account for OTU oversplitting. The final OTU table contained 41 334 OTUs. OTUs were taxonomically annotated using SINTAX (Edgar, 2016), as implemented in VSEARCH, against the BOLDistilled database (July 2025 version; Rognes et al., 2016).

### Statistical analyses

Data exploration and analyses were conducted in R v. 4.5.2 (R Core Team, 2025). Rarefaction curves were generated to evaluate sequencing coverage across samples (Appendix figure 1). Due to relatively low sequencing depth, no normalization or rarefaction was done. Instead, log-transformed sequencing depth was included as a covariate in all models to account for variation in read numbers among samples. Figures were generated using the ggplot2 package (Gómez Rubio, 2017), and alpha diversity metrics were estimated with phyloseq (McMurdie and Holmes, 2013).

To test whether arthropod diversity changed across the successional gradient or forest management (H1, H3), alpha diversity was quantified as OTU richness and Pielou’s evenness. Richness was analyzed using generalized linear mixed-effects models (GLMM) with a negative binomial error distribution and log link function implemented in the glmmTMB package (Brooks et al., 2017). Evenness was analyzed using linear mixed-effects models (LMM) for deadwood in the lmer package ((Bates et al., 2015). Site ID was included as a random intercept to account for non-independence between samples from the same site. For organic soil layers, evenness was analyzed using a linear model because the site-level random effect was estimated to zero.

The models included forest management type, decay stage (DW1-DW4/5), log-transformed sequencing depth, pH, C:N ratio, soil volume, slope, and mean annual temperature (MAT) as fixed effects, depending on substrate. Prior to model selection, multicollinearity among explanatory variables was checked using variance inflation factors (all < 3). Final models were decided using backward model selection based on Akaike’s Information Criterion (AIC). Variables directly related to the study hypotheses (decay stage, forest management type) and sequencing depth were retained during model selection. Model assumptions were evaluated using simulated residual diagnostics in the package DHARMa (Hartig, 2026). To evaluate successional patterns across the complete deadwood–soil gradient, an additional set of models was fitted using a five-level substrate factor with DW1, DW2, DW3, DW4/5, and organic soil layer. Residual diagnostics indicated no major deviations from model assumptions, including no evidence of substantial overdispersion, heteroscedasticity, or influential outliers.

To investigate if arthropod communities were shaped by succession (H2), community composition in deadwood and organic soil layer was explored using ordination and permutation-based multivariate analyses. First, community composition patterns were visualized with non-metric multidimensional scaling (NMDS) implemented in the vegan package (Oksanen et al., 2015). OTU abundance tables were transformed to relative abundances and Hellinger-transformed prior to ordination analysis, and Bray-Curtis dissimilarities were calculated from the transformed data. Samples with fewer than 100 reads and OTUs occurring in fewer than two samples were removed prior to multivariate analyses, resulting in the removal of 219 samples and 447 OTUs. The majority of the removed OTUs were extremely rare, and most were only present in one sample. In addition, eight outliers were removed from the ordination space before the final ordination analyses. This was done to improve the interpretability of the ordination and reduce distortion of community patterns. As a result, the ordination included 262 samples and 367 OTUs.

Secondly, variation in community composition among substrates, decay stages, and forest types was analyzed separately for the deadwood dataset and the combined deadwood-organic soil dataset using permutational multivariate analysis of variance (PERMANOVA) from the adonis2 function in vegan (Oksanen et al., 2015). Initial PERMANOVA models included interaction terms between forest type and decay stage (deadwood dataset) or substrate group (combined deadwood–soil dataset). As these interactions were not significant, they were omitted from the final models. We restricted permutations within sites to account for the nested sampling design, and evaluated the significance with 9 999 permutations. The betadisper function was used for assessing the homogeneity of multivariate dispersion between groups. Finally, constrained analyses of principal coordinates (CAP) were used to show differences in community composition along the successional gradient. Correlations between environmental variables and community composition were assessed by fitting environmental variables to the NMDS ordination using the envfit function. For all model- and permutation-based analyses, statistical significance was evaluated at α = 0.05.

We further explored co-occurrence patterns between arthropod OTUs and substrate types with a network-based analysis. Here, co-occurrences refer to OTUs that were significantly associated with individual substrate groups or combinations of substrate groups. This allows us to assess the extent to which arthropod OTUs were shared across the decay gradient and the organic soil layers. Co-occurrences associated with the decay stages and organic soil layers were identified using the multi-level pattern analysis in the package indicspecies (Cáceres and Legendre, 2009). This identified OTUs significantly associated with specific or combined substrate groups based on indicator values calculated from species occurrence and abundance patterns (Cáceres and Legendre, 2009). Indicator significance was tested by 9 999 permutations, and Benjamini-Hochberg adjusted p-values were used to account for multiple testing with α = 0.05 (Benjamini and Hochberg, 1995).

## Results

By use of DNA metabarcoding we detected 741 arthropod OTUs across the analyzed deadwood and soil samples. In total, 565 OTUs were found in deadwood (445 samples) and 291 OTUs in soil (44 samples), while 115 OTUs were shared between substrates. Of the OTUs identified to class level, insects represented the largest group (268 OTUs). Previously clear-cut (CC) and near-natural (NN) forests hosted comparable numbers of OTUs (511 and 541, respectively), but only 311 OTUs were shared between forest types.

### Diversity

In deadwood, decay stage had a significant effect on arthropod OTU richness (χ² = 11.29, df = 3, p = 0.010), with an overall positive trend across the decay gradient (Figure 2). From decay stage 1 to 4, predicted richness increased from 7.39 (95% CI: 6.47–8.46) to 10.15 (8.59– 12.00) OTUs per sample. In addition, richness was positively related to sequencing depth (estimate = 0.126 ± 0.025 SE, p < 0.001) and moisture (estimate = 0.0045 ± 0.0021 SE, p = 0.032), whereas forest management type showed no effect (p = 0.269). Despite a visual trend, evenness was not affected by decay stage or forest type (all p > 0.1; Figure 2). However, evenness decreased strongly with sequencing depth (estimate = −0.105 ± 0.012 SE, p < 0.001).

**Figure 2.**
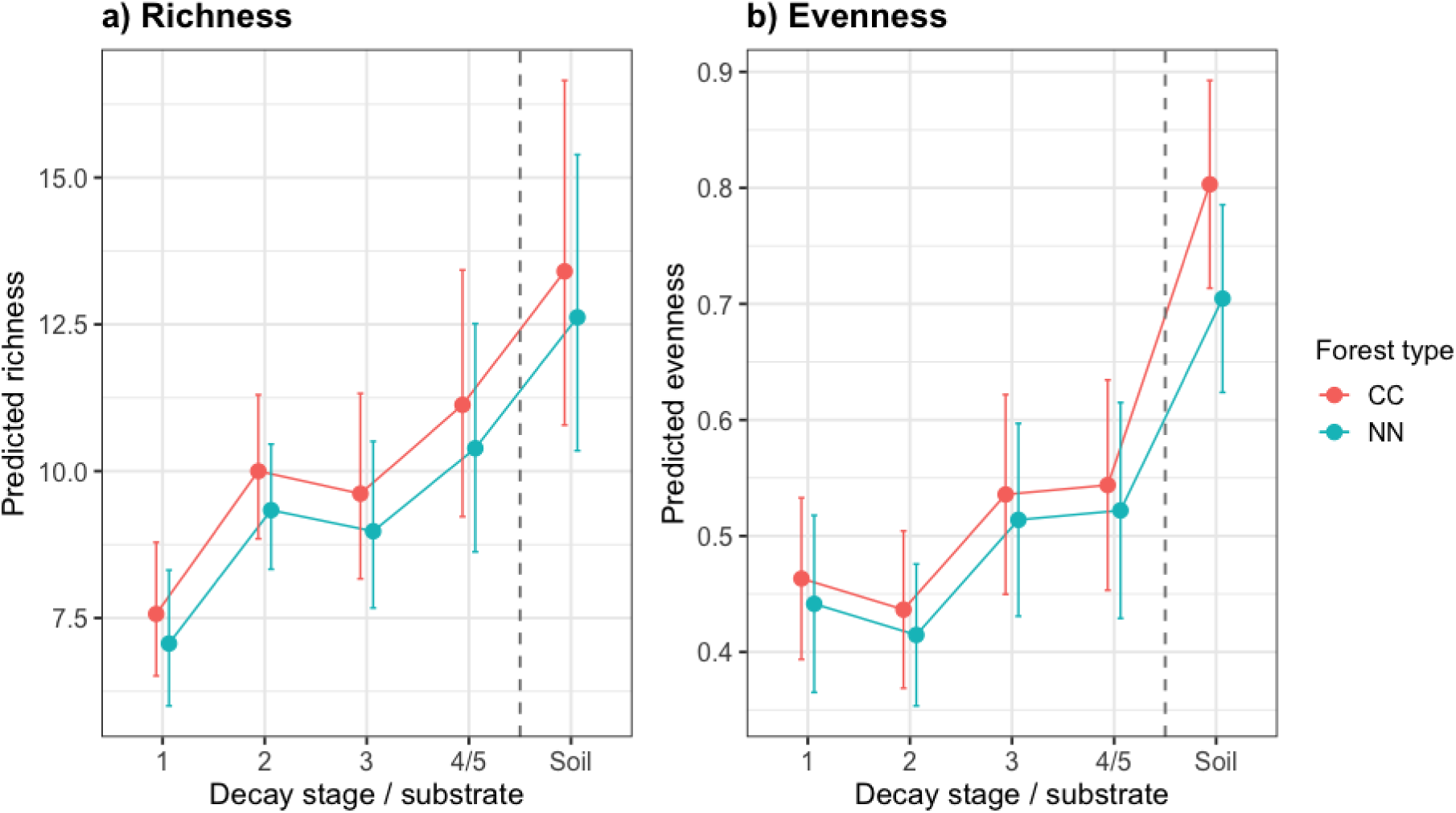
Model-predicted arthropod OTU richness (a) and evenness (b) across deadwood decay stages (DW1–4/5) and organic soil layers in previously clear-cut (CC) and near-natural (NN) forests. Points represent predicted values from generalized mixed models for deadwood samples and separate models for soil samples, while error bars show 95% confidence intervals. The dashed vertical line separate deadwood from organic soil layers.

For organic soil layers, richness increased significantly with sequencing depth (estimate = 0.506 ± 0.032 SE, p < 0.001) and pH (estimate = 0.582 ± 0.098 SE, p < 0.001), and decreased with soil bulk density (estimate = −0.0028 ± 0.0013 SE, p = 0.039). Forest type had no significant effect on richness (p = 0.732). Also, neither forest type nor sequencing depth significantly influenced arthropod evenness (forest type: estimate = −0.085 ± 0.059 SE, p = 0.154; sequencing depth: estimate = 0.012 ± 0.017 SE, p = 0.511).

When deadwood decay stages and organic soil layers were analyzed together, arthropod richness differed significantly among substrate groups (χ² = 74.62, df = 4, p < 0.001; Figure 2). Pairwise comparisons showed that organic soil layers supported significantly higher richness than all deadwood decay stages, while richness differences among DW2–DW4/5 were relatively small and mostly non-significant. Arthropod evenness also differed significantly among substrate groups (χ² = 26.66, df = 4, p < 0.001), with organic soil layers showing significantly higher evenness than all deadwood decay stages. In contrast, evenness differences among deadwood decay stages were small and non-significant (all p > 0.05). The interaction between substrate and forest type was not significant for evenness (X^2^ = 8.68, df = 4, p = 0.070).

### Community composition

Different successional stages hosted distinct arthropod communities (Figure 3). In deadwood, Insecta was the most abundant class but were less prominent in later decay stages (DW3 and DW4/5). Curculionidae (true weevils) was the most abundant family overall Appendix S1, Table S2), with particularly high dominance in DW1. In general, Arachnida were more abundant in DW2–DW4/5 and soil, mostly represented by oribatid families such as Carabodidae in deadwood and Nanhermanniidae in organic soil layers. Collembola (springtails) were the most common class in soil, but were also present in all deadwood substrates, although with low relative abundance in DW2. Some families differed in abundances between previously clear-cut and near-natural forests. For instance, Sciaridae (dark-winged fungus gnats) was more abundant in previously clear-cut forests (CC), while Ptinidae (wood-borer beetles) was more abundant in near-natural forests (NN).

**Figure 3.**
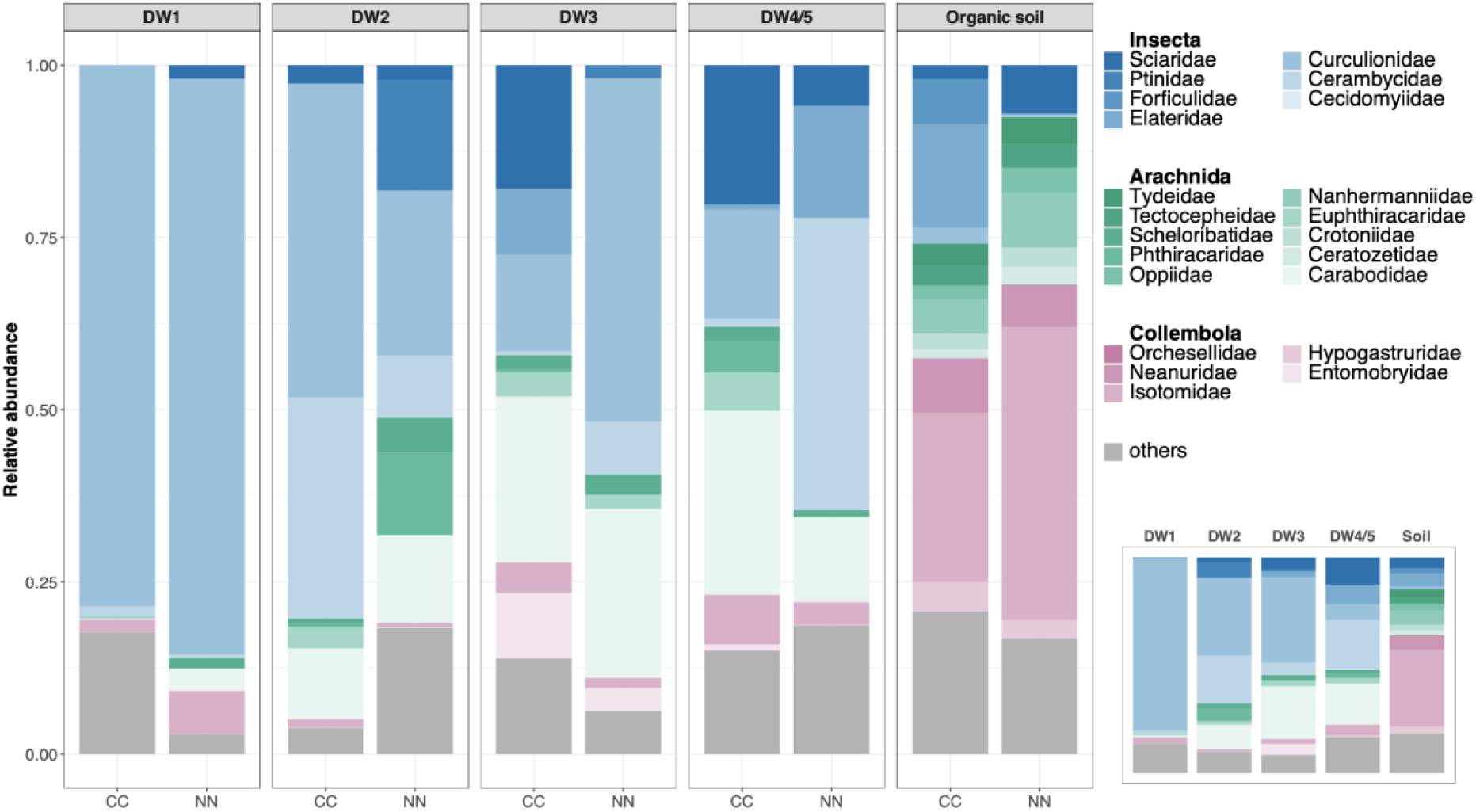
Relative sequence abundance of arthropod families across deadwood decay stages (DW1–DW4/5) and organic soil layers in previously clear-cut (CC) and near-natural (NN) forests. Colour palettes indicate major arthropod classes, with blue shades representing Insecta, green shades Arachnida, pink shades Collembola, and grey representing both unknown OTUs and all families with mean relative abundance below 1% across forest types (“others”). Relative abundances are based on proportional sequence reads within each substrate and forest type combination. The inset panel shows relative sequence abundances pooled across forest types for each decay stage and organic soil layers.

Ordination (NMDS and CAP) showed clear structuring of arthropod communities along the decay gradient, with early decay stages separated from intermediate and late stages (Figure 4; Appendix S1, Figure 1). Consistent with these patterns, PERMANOVA restricted to deadwood samples showed that community composition differed significantly among decay stages *(F* = 1.83, R² = 0.024, p = 0.001). Relative moisture *(F* = 3.07, R² = 0.014, p = 0.001), log-transformed volume *(F* = 2.36, R² = 0.010, p = 0.007), pH *(F* = 2.07, R² = 0.009, p = 0.019), and C:N ratio *(F* = 2.02, R² = 0.009, p = 0.008) were also significant predictors of community composition. When accounting for these variables, forest type (clear-cut vs near-natural) did not significantly affect arthropod community composition (*F* = 1.17, R² = 0.005, p = 0.198). Homogeneity of multivariate dispersion did not differ between decay stages (betadisper: *F* = 0.42, p = 0.74).

**Figure 4.**
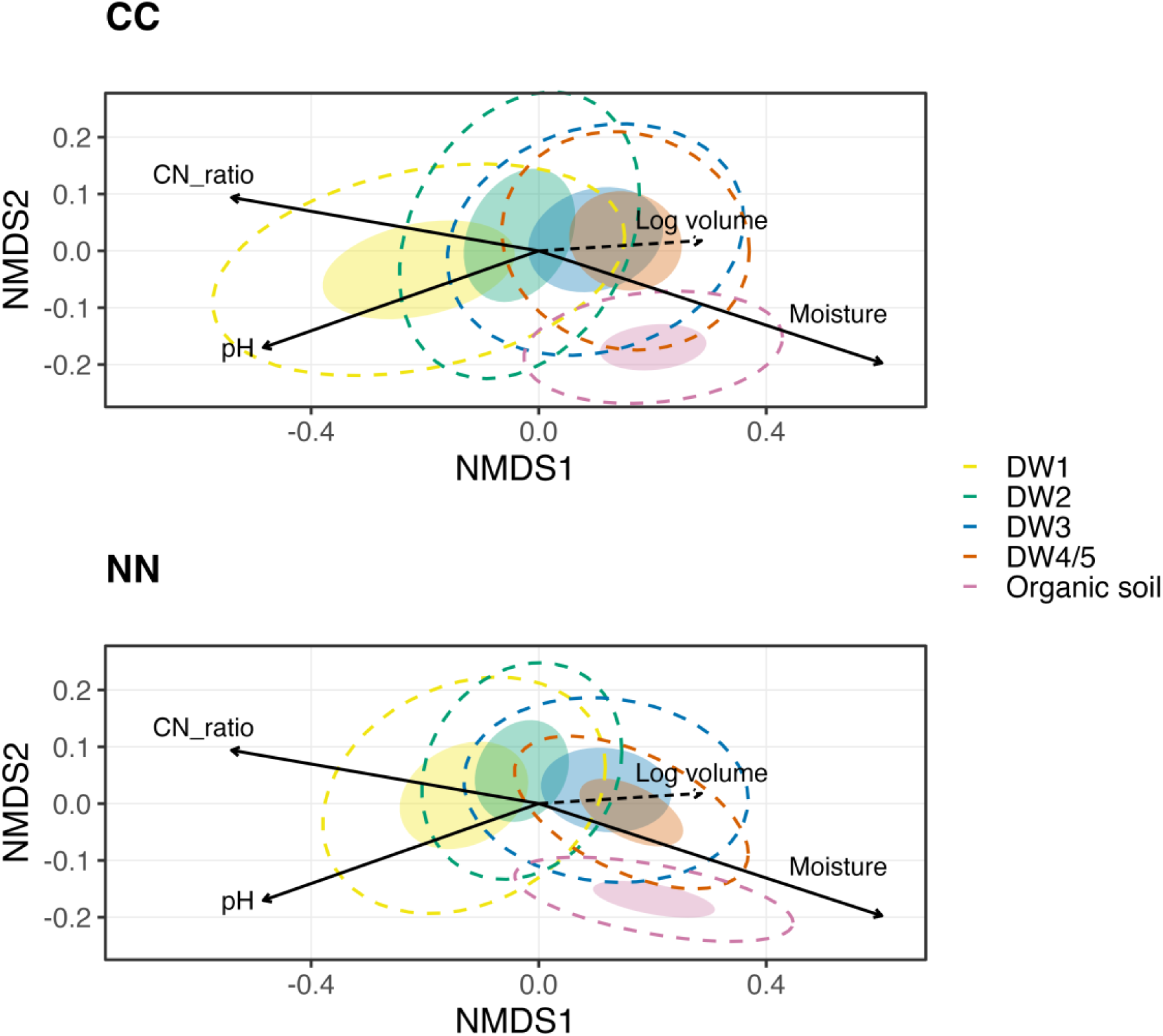
Non-metric multidimensional scaling (NMDS) ordination of arthropod community composition of previously clear-cut (CC) and near-natural (NN) forests based on Bray–Curtis dissimilarities across deadwood decay stages (DW1–DW4/5) and organic soil layers. Filled ellipses show 50% confidence regions around group centroids, while dashed ellipses show 95% confidence regions. Arrows indicate significant environmental gradients fitted using envfit (p ≤ 0.05), with arrow length proportional to the correlation strength. Except for log volume (dashed line), all variables were measured for both deadwood and organic soil layers. Stress = 0.174.

In a PERMANOVA including both deadwood and soil samples, community composition was significantly different between deadwood and organic soil layers *(F* = 7.28, R² = 0.099, p < 0.001). Organic soil layers formed a distinct cluster that was more similar to late decay than early decay deadwood (Figure 4). In the NMDS, plot centroids of communities from previously clear-cut forests were consistently at more extreme positions, lying further away from the overall centroid than those from the corresponding near-natural forests (Appendix S1, Figure S2). Consistent with this pattern, multivariate dispersion was significantly higher in previously clear-cut than in near-natural forests (betadisper: *F* = 4.00, p = 0.045).

### Co-occurrence patterns

We identified in total 85 arthropod OTUs with significant associations to specific deadwood decay stages and/or soil (‘co-occurrences’; Figure 5; Appendix S1, Table S1). Among these, 65 (76.5%) were significantly associated with one specific decay stage (17) or organic soil layers (48). Some OTUs displayed broader substrate affinities, with 12 OTUs (14.1%) associated with two substrates, and eight (9.4%) associated with three or four substrate types. More OTUs were shared between DW4/5 and organic soil layers (7) than between earlier decay stages and organic soil layers (4) (Figure 5). In addition, co-occurrence patterns differed somewhat between forest types. Previously clear-cut and near-natural forests supported partially distinct co-occurrence patterns, with 217 and 244 arthropod OTUs with significant associations, respectively. In near-natural forests, several arthropod OTUs co-occurred with both DW4/5 and organic soil layers, whereas no such co-occurrences were identified in previously clear-cut forests (Figure 5; Appendix S1, Table S1). Additionally, near-natural forests had more arthropod OTUs with significant associations to organic soil layers than was found in previously clear-cut forests (29 vs. 21), but fewer co-occurrences with DW1 (1 vs. 4).

**Figure 5.**
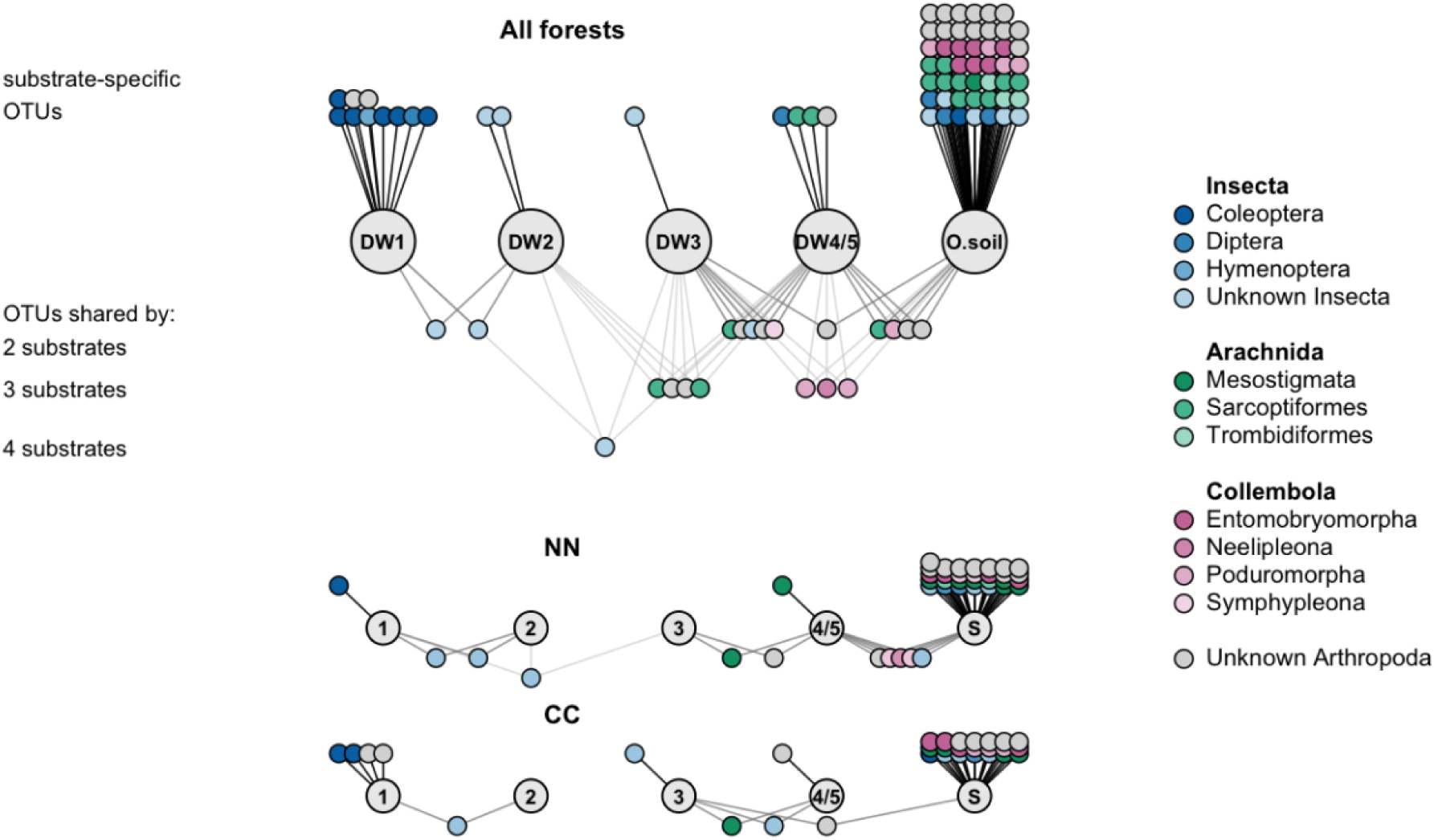
Tripartite network based on indicator analysis showing significant associations between 85 OTUs and larger grey circles show deadwood (DW1–4/5) and organic soil substrate groups. Circles at the top represent OTUs significantly associated with only one substrate type, while circles underneath show OTUs linked to 2-4 substrate types. OTUs are colored by order, with those belonging to class Insecta shown in blue shades, Arachnida in green, Collembola in pink, and unidentified arthropods in grey. The top figure shows co-occurrences for both forest types, while those only associated with near-natural or previously clear-cut forests are displayed in the bottom figures.

## Discussion

Our study demonstrates that arthropod succession in Norway spruce deadwood extends further than the deadwood itself, with communities gradually converging towards those in organic soil layers during decay. As expected, arthropod richness increased along the deadwood decay gradient, while community composition changed as predicted from early to late decay deadwood, with late-decay communities becoming increasingly similar to those in organic soil layers. Community overlap between deadwood and organic soil layers also increased during later stages of decay, supporting the idea that highly decayed logs function as transitional habitats linking these two major forest substrates. Although forest management had limited effects on overall diversity patterns, previously clear-cut and near-natural forests still supported partially distinct arthropod communities.

### Succession in deadwood

Arthropod communities showed a clear successional trajectory during deadwood decay, characterized by increasing species richness and gradual changes in community composition from early to late decay stages. Richness increased along the decay gradient, which is consistent with the habitat-heterogeneity hypothesis (MacArthur and MacArthur, 1961, Stein et al., 2014). The higher diversity of available microhabitats likely enables arthropods with different feeding strategies and habitat requirements to coexist, contributing to the higher richness observed in later decay stages (Ulyshen, 2016). In contrast to richness, arthropod evenness did not change significantly along the decay gradient. This indicates that even though an increasingly higher number of taxa were detected during decay, the relative distribution of abundances among taxa remained similar in all decay stages.

Relative moisture content was positively correlated with arthropod richness and community composition in deadwood. Moisture is a key element for fungal growth in deadwood (Brischke and Alfredsen, 2020, Gómez-Brandón et al., 2020). Consequently, its increase likely affects arthropod communities indirectly by increasing food availability for fungivorous and detritivorous arthropods (Boddy, 2001, Rajala et al., 2012, Seibold et al., 2016). This suggests that moisture may have partly indirect effects on richness and community composition through increased fungal growth. This is consistent with succession studies from other ecosystems where environmental changes correlated to advancing succession act as filters that shape community composition over time (Walker and del Moral, 2003, Fukami, 2015).

Targeting the broader arthropod community with COI metabarcoding, the overall increase in richness reflects the cumulative responses of multiple arthropod groups with different successional trajectories. In addition to the well-known deadwood-associated beetles (Stokland et al., 2012), our metabarcoding data detected other insects, oribatid mites and springtails often overlooked in previous studies. Increased richness with advancing decay was also found in the only other study we are aware of focusing on arthropods in conifer deadwood (Kim et al., 2013). In contrast, studies focusing exclusively of beetles have found different successional patterns. Early-decay deadwood supports numerous bark- and phloem-associated beetles (Ulyshen and Hanula, 2010), and a recent twelve-year study of Norway spruce reported a U-shaped richness pattern for beetles, with richness peaking both in early and late decay (Lettenmaier et al., 2026). Despite similarities with our findings for relative abundance in early decay, our results likely reflect the arrival of a large variety of taxa after the initial colonization of bark beetles. For instance, other prevalent taxa from our study, such as true flies and oribatid mites, have been found to increase in richness with decay stage (Hövemeyer and Schauermann, 2003, Skubała and Sokołowska, 2006). This likely contributed to the overall increase in richness found in our data. Limitations in reference databases and potential primer bias likely influenced taxonomic resolution and detection probabilities for some arthropod groups (Elbrecht et al., 2019, Ficetola et al., 2025). However, the arthropods detected in different decay stages and organic soil layers are all ecologically meaningful and consistent with previous research based on morphology (Stokland et al., 2012, Parisi et al., 2018).

As hypothesised, arthropod community composition changes predictably along the decay gradient. This turnover is consistent with previous studies showing that deadwood communities are structured by changes in substrate conditions during decay (Lepinay et al., 2021, Seibold et al., 2023). In addition to influencing richness, moisture also showed the strongest relationship with arthropod composition in the ordination analyses. On the other hand, C:N ratio and pH only had significant effects on community composition, not richness. pH and C:N ratio likely influence fungal growth (Rajala et al., 2012), which indirectly shapes arthropod community composition because fungi is an important food resource for many deadwood-associated arthropods (Stokland et al., 2012). These findings suggest that the environmental changes occurring during decay contribute to the observed turnover of arthropod communities.

### From deadwood to soil

Deadwood and organic soil layers are both two of the major forest substrates, but are fundamentally different habitats. Deadwood is discrete, spatially isolated and ephemeral, whereas organic soil layers form a continuous and relatively persistent habitat. Our results, however, show that deadwood succession extends beyond the deadwood itself and continues into the forest floor, representing a gradual ecological succession from deadwood to organic soil layers. The community composition of arthropods in the organic soil layers was most similar to the communities in late-decay deadwood, indicating convergence between these habitats during advanced decay. Early decay stages were dominated by specialist bark beetles, whereas those from later decay stages contained a broader range of fungivorous, detritivorous, and soil-associated taxa, including oribatid mites and springtails. In addition, later decay stages shared multiple co-occurrences with communities in organic soil layers. Early decay deadwood remained characterized by specialists such as bark beetles, and few co-occurrences were specific to DW2 and DW3, while springtails and oribatid mites were associated with late decay deadwood. This suggests that late decay deadwood is suitable for colonization by soil-associated arthropods, likely reflecting increasingly soil-like conditions in highly decomposed wood (Mäkipää et al., 2017, Parisi et al., 2018).Together, these findings suggest gradual community turnover throughout decay and indicate that later decay stages may function as transitional habitats between communities associated with deadwood and organic soil layers. Rather than representing the endpoint of deadwood succession, late decay logs seem to function as ecological transition zones that connect two major forest substrates.

Although organic soil layers supported significantly higher arthropod richness than all deadwood decay stages, this was partly expected due to differences in sampling methodology, as soil samples were pooled samples from a larger part of the plot. Nevertheless, similar convergence trends, where deadwood is increasingly colonized by soil-associated fungi, have been described between fungal communities in Norway spruce logs and soil (Rajala et al., 2012, Mäkipää et al., 2017). This suggests that arthropod succession follows the same patterns, potentially mediated by changes in fungal communities and the resources they provide during decay.

### Importance of forest management

Forest management had limited effects on overall arthropod richness and community composition, contrasting our original hypothesis. Nevertheless, arthropod communities in previously clear-cut and near-natural forests occupied different positions in ordination space, suggesting subtle differences in community composition. Although these differences were not statistically significant, they could reflect differences in species pools caused by overall habitat qualities; deadwood volume was on average 3.6 times higher in our near-natural forests than in previously clear-cut forests (Asplund et al., 2024), potentially supporting larger arthropod populations and increasing colonization probability of deadwood (Lassauce et al., 2011, Kraut et al., 2016, Haeler et al., 2021). Many deadwood-associated arthropods have limited dispersal abilities (Stokland et al., 2012, Seibold et al., 2015) and may be sensitive to the lower habitat availability in previously clear-cut forests. The co-occurrence networks further suggest that there is a stronger overlap between communities in late decay deadwood and organic soil layers in near-natural than in previously clear-cut forests. This pattern likely reflects the differences in volume of sampled deadwood logs, where logs from near-natural forests had 46% higher volume of late decay stages than in previously clear-cut forests. The near-natural forests could therefore maintain a stronger ecological connectivity between deadwood and soil, with possible implications for wood decay, soil processes and habitat use by arthropods otherwise confined to organic soil layers (Fujii et al., 2023). These findings highlight the importance of a continuous supply of deadwood logs to maintain species and functions related to the gradual transition between deadwood and soil.

### Concluding remarks

Our study demonstrates increasing species richness and a clear turnover in arthropod composition across the deadwood decay gradient and into organic soil layers. The communities shifted predictably from beetle specialists to fungivorous, detritivorous, and soil-associated taxa such as true flies, mites and springtails. We also found that highly decayed deadwood functions as a transitional habitat between deadwood and forest soil, a transition that may be broken by clear-cut forestry. Maintaining a continuous supply of deadwood across all decay stages is therefore likely to help preserve this ecological transition in managed forests. Beyond their implications for forest management, our results demonstrate how ephemeral habitats can serve as model systems for studying general mechanisms of ecological succession. As deadwood decays, increasing habitat heterogeneity, changing environmental conditions, and shifts in resource availability drive predictable community changes like successional dynamics described across a wide range of successional ecosystems. Finally, our study shows that DNA metabarcoding can be a powerful tool for studying arthropods in deadwood, but future studies should include both bark and wood samples to better capture to better capture communities associated with the earliest stages of decay.

## Acknowledgements

The Research Council of Norway is acknowledged for financial support to the EcoForest project (No. 320722). Research permits for nature reserves were issued by the County Governor of Oslo and Viken (2021/22482) and the County Governor of Innlandet (2021/7972). We acknowledge forest owners for allowing research on their properties.

Rebecca Biong, Juliette Combret, Thibaut Sciaccitano, and Anders Kvalvåg Wollan are acknowledged for setting up the EcoForest plots and for data collection. Thank you to Tom Hellik Hoften (Biofokus), Siri Khalsa (Biofokus), and Erik Möller for deadwood surveys. The bioinformatics analyses were performed on Saga, provided by Sigma2, the National Infrastructure for High-Performance Computing and Data Storage in Norway.

