## Appendix S1 for "Arthropod succession from deadwood to soil: increasing diversity and transitions between major forest substrates"

Supplementary Information for

Arthropod succession from deadwood to soil:

increasing diversity and transitions between major forest substrates

William Esbjug Gromstad

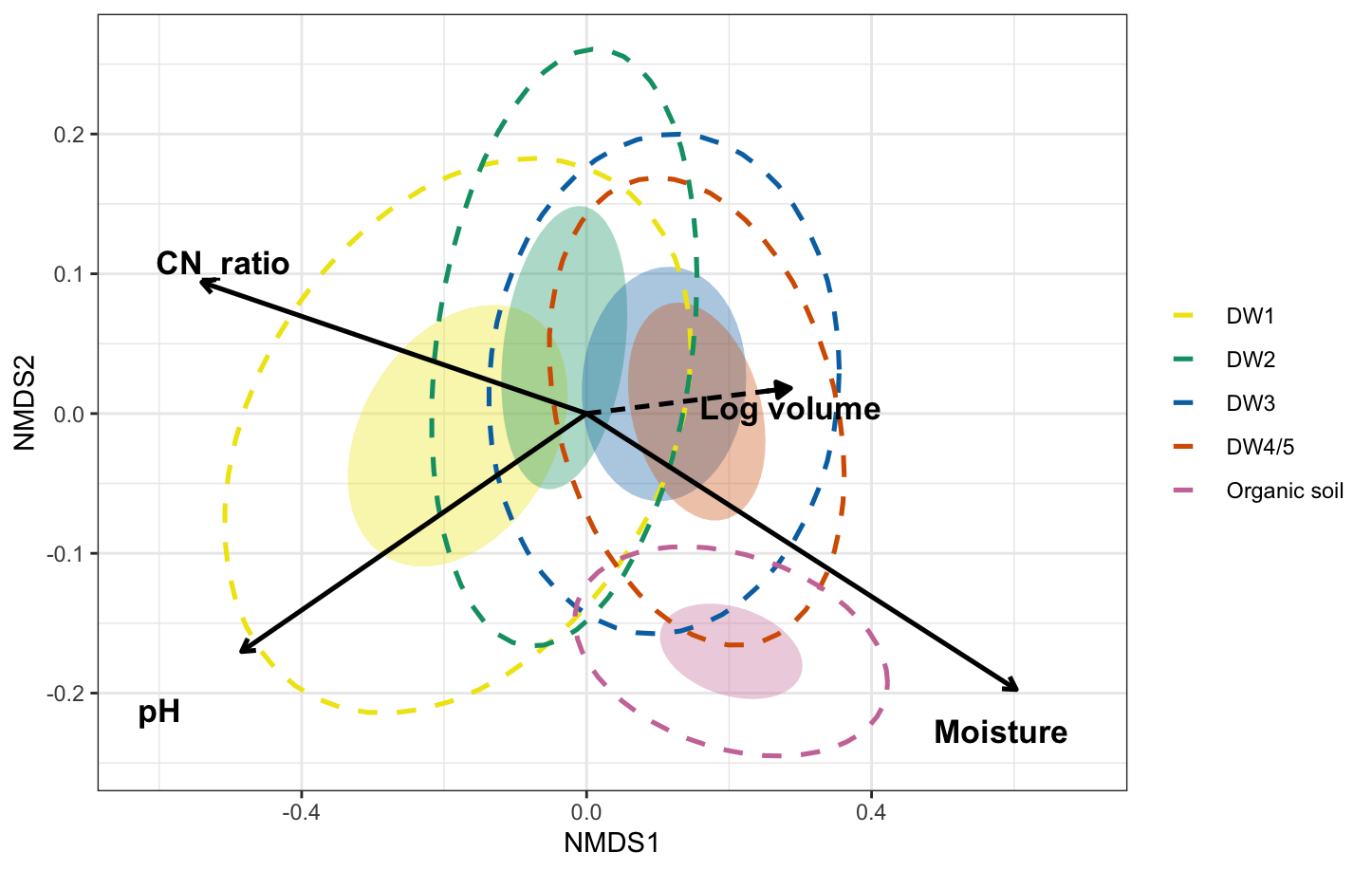

Figure S1: Canonical analysis of principal coordinates (CAP) showing arthropod community composition across deadwood decay stages (DW1–DW4/5) and organic soil layersbased on Bray–Curtis dissimilarities. Filled ellipses represent 50% group dispersion and dashed ellipses represent 95% group dispersion around group centroids. Arrows indicate significant environmental variables fitted onto the ordination using envfit, including moisture (%), C:N ratio, pH, and log volume for deadwood samples.

Community composition in organic soil layers was analysed together with deadwood decay stages, also using PERMANOVA based on Bray–Curtis dissimilarities. Composition was significantly different among decay stages and organic soil layers (F = 7.28, R² = 0.099, p < 0.001). Pairwise comparisons showed that communities in organic soil layers differed significantly from all deadwood decay stages (all p < 0.001), while all decay stages differed significantly from each other except for the two most advanced stages (DW3 vs DW4/5; p = 0.16), indicating convergence of community composition in late decay.

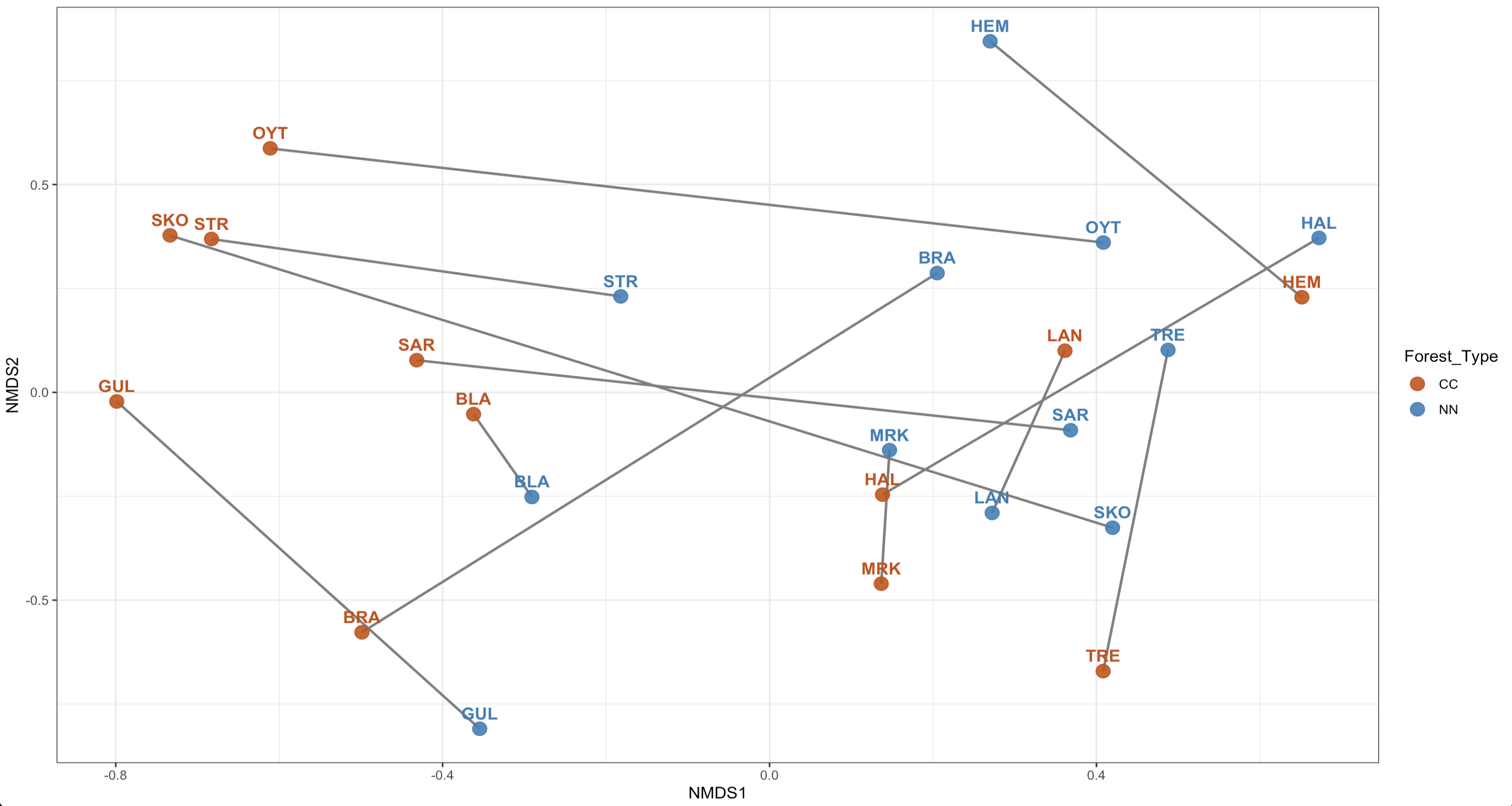

Figure S2: Non-metric multidimensional scaling (NMDS) ordination showing differences in arthropod community composition between previously clear-cut (CC) and near-natural (NN) forests within site pairs based on Bray–Curtis dissimilarities. Points represent individual plots and labels indicate site identities. Lines connect paired CC and NN plots from the same site, illustrating within-pair compositional differences between forest types.

Table S1: Arthropod OTUs with significant associations to individual or combined deadwood decay stages and organic soil layers in all forests, only previously clear-cut forests, and only near-natural forests, identified by multi-level pattern analysis (indicator species analysis). OTUs significantly associated with one or more substrate groups (Benjamini–Hochberg adjusted P < 0.05) are shown together with their associated substrate(s), indicator values (IndVal), and adjusted P-values.

### All forests

| Association | Indicator value | Adjusted p | Class | Order | Family | Genus | Species |
| --- | --- | --- | --- | --- | --- | --- | --- |
| DW1 | 0.567 | 0.0011 | Insecta | Coleoptera | Curculionidae | Pityogenes | Pityogenes_chalcographus |
| DW1 | 0.477 | 0.0011 | Insecta | Coleoptera | Curculionidae | Pityogenes | Pityogenes_chalcographus |
| DW1 | 0.42 | 0.0019 | — | — | — | — | — |
| DW1 | 0.409 | 0.0163 | Insecta | Hymenoptera | — | — | — |
| DW1 | 0.367 | 0.0072 | — | — | — | — | — |
| DW1 | 0.304 | 0.0329 | Insecta | Coleoptera | Curculionidae | Polygraphus | Polygraphus_punctifrons |
| DW1 | 0.304 | 0.0132 | Insecta | Coleoptera | Curculionidae | Pityogenes | Pityogenes_chalcographus |
| DW1 | 0.302 | 0.0451 | Insecta | Diptera | Pallopteridae | Toxonevra | Toxonevra_usta |
| DW1 | 0.301 | 0.0208 | Insecta | Coleoptera | Curculionidae | Pityogenes | Pityogenes_chalcographus |
| DW1 | 0.299 | 0.0465 | — | — | — | — | — |
| DW1 | 0.277 | 0.0233 | Insecta | Coleoptera | Curculionidae | Pityogenes | Pityogenes_chalcographus |
| DW1+DW2 | 0.58 | 0.0011 | Insecta | — | — | — | — |
| DW1+DW2 | 0.452 | 0.0051 | Insecta | — | — | — | — |
| DW1+DW2 | 0.349 | 0.0472 | Insecta | — | — | — | — |
| DW1+DW2+DW3+DW4/5 | 0.727 | 0.0011 | Insecta | — | — | — | — |
| DW2 | 0.5 | 0.0036 | Insecta | — | — | — | — |
| DW2 | 0.365 | 0.0384 | Insecta | — | — | — | — |
| DW2+DW3+DW4/5 | 0.532 | 0.0061 | Arachnida | Sarcoptiformes | Carabodidae | Carabodes | Carabodes_areolatus |
| DW2+DW3+DW4/5 | 0.442 | 0.0047 | — | — | — | — | — |
| DW2+DW3+DW4/5 | 0.368 | 0.0329 | — | — | — | — | — |
| DW2+DW3+DW4/5 | 0.367 | 0.0367 | Arachnida | Sarcoptiformes | Scheloribatidae | Scheloribates | Scheloribates_pallidulus |
| DW3 | 0.557 | 0.0011 | Insecta | — | — | — | — |
| DW3+DW4/5 | 0.642 | 0.0011 | Arachnida | Sarcoptiformes | Carabodidae | Carabodes | — |
| DW3+DW4/5 | 0.523 | 0.0011 | — | — | — | — | — |
| DW3+DW4/5 | 0.41 | 0.0011 | Insecta | — | — | — | — |
| DW3+DW4/5 | 0.334 | 0.0169 | — | — | — | — | — |
| DW3+DW4/5 | 0.284 | 0.0277 | Collembola | Symphypleona | Sminthuridae | Lipothrix | Lipothrix_lubbocki |
| DW3+DW4/5+Organic | 0.474 | 0.0019 | Collembola | Poduromorpha | Hypogastruridae | Willemia | Willemia_anophthalma |
| DW3+DW4/5+Organic | 0.403 | 0.0047 | Collembola | Neelipleona | Neelidae | Megalothorax | — |
| DW3+DW4/5+Organic | 0.363 | 0.0148 | Collembola | Poduromorpha | Neanuridae | Neanura | Neanura_muscorum |
| DW3+Organic | 0.462 | 0.0027 | — | — | — | — | — |
| DW4/5 | 0.391 | 0.0011 | Arachnida | Sarcoptiformes | Oribotritiidae | Protoribotritia | — |
| DW4/5 | 0.363 | 0.0276 | — | — | — | — | — |
| DW4/5 | 0.345 | 0.0047 | Insecta | Diptera | Sciaridae | Pnyxia | — |
| DW4/5 | 0.277 | 0.0166 | Arachnida | Sarcoptiformes | Caleremaeidae | Caleremaeus | Caleremaeus_monilipes |
| DW4/5+Organic | 0.333 | 0.0051 | Arachnida | Sarcoptiformes | Nothridae | Nothrus | Nothrus_silvestris |
| DW4/5+Organic | 0.314 | 0.0129 | Collembola | Poduromorpha | Neanuridae | Micranurida | Micranurida_forsslundi |
| DW4/5+Organic | 0.294 | 0.0342 | — | — | — | — | — |
| DW4/5+Organic | 0.272 | 0.0347 | — | — | — | — | — |
| Organic | 0.814 | 0.0011 | Collembola | Entomobryomorpha | Isotomidae | Isotomiella | Isotomiella_minor |
| Organic | 0.74 | 0.0011 | — | — | — | — | — |
| Organic | 0.707 | 0.0011 | Arachnida | Sarcoptiformes | Nanhermanniidae | Nanhermannia | Nanhermannia_sellnicki |
| Organic | 0.584 | 0.0011 | — | — | — | — | — |
| Organic | 0.538 | 0.0011 | Insecta | — | — | — | — |
| Organic | 0.512 | 0.0011 | Collembola | Entomobryomorpha | Isotomidae | Folsomia | Folsomia_sensibilis |
| Organic | 0.512 | 0.0011 | — | — | — | — | — |
| Organic | 0.488 | 0.0011 | Insecta | Diptera | Sciaridae | — | — |
| Organic | 0.488 | 0.0011 | Arachnida | Sarcoptiformes | Ceratozetidae | Ceratozetella | Ceratozetella_thienemanni |
| Organic | 0.463 | 0.0011 | Arachnida | Sarcoptiformes | Tectocepheidae | Tectocepheus | Tectocepheus_velatus |
| Organic | 0.463 | 0.0011 | Collembola | Entomobryomorpha | Isotomidae | Isotomiella | Isotomiella_minor |
| Organic | 0.448 | 0.0011 | — | — | — | — | — |
| Organic | 0.436 | 0.0011 | Insecta | Coleoptera | Elateridae | Athous | Athous_subfuscus |
| Organic | 0.436 | 0.0011 | Collembola | Poduromorpha | Neanuridae | Anurida | Anurida_granulata |
| Organic | 0.436 | 0.0011 | Arachnida | Trombidiformes | Tydeidae | — | — |
| Organic | 0.417 | 0.0011 | Collembola | Poduromorpha | Onychiuridae | Protaphorura | Protaphorura_pseudovanderdrifti |
| Organic | 0.412 | 0.0011 | Collembola | Poduromorpha | Neanuridae | Friesea | Friesea_mirabilis |
| Organic | 0.408 | 0.0011 | Arachnida | Sarcoptiformes | Oppiidae | Oppiella | Oppiella_nova |
| Organic | 0.408 | 0.0011 | — | — | — | — | — |
| Organic | 0.403 | 0.0019 | Insecta | — | — | — | — |
| Organic | 0.4 | 0.0011 | — | — | — | — | — |
| Organic | 0.394 | 0.0019 | Arachnida | Trombidiformes | Eupodidae | — | — |
| Organic | 0.378 | 0.0011 | — | — | — | — | — |
| Organic | 0.378 | 0.0011 | Insecta | Diptera | Bibionidae | Bibio | — |
| Organic | 0.367 | 0.0072 | — | — | — | — | — |
| Organic | 0.345 | 0.0019 | Collembola | Entomobryomorpha | Isotomidae | Parisotoma | Parisotoma_notabilis |
| Organic | 0.345 | 0.0011 | Arachnida | Sarcoptiformes | Oppiidae | Oppiella | Oppiella_nova |
| Organic | 0.345 | 0.0011 | Insecta | — | — | — | — |
| Organic | 0.341 | 0.01 | — | — | — | — | — |
| Organic | 0.328 | 0.0077 | Arachnida | Sarcoptiformes | Crotoniidae | Platynothrus | Platynothrus_peltifer |
| Organic | 0.314 | 0.0407 | Insecta | — | — | — | — |
| Organic | 0.309 | 0.0057 | Insecta | Diptera | Sciaridae | Corynoptera | Corynoptera_boletiphaga |
| Organic | 0.309 | 0.0043 | Collembola | Entomobryomorpha | Isotomidae | — | — |
| Organic | 0.309 | 0.0051 | Collembola | Poduromorpha | Hypogastruridae | Willemia | Willemia_anophthalma |
| Organic | 0.309 | 0.0051 | — | — | — | — | — |
| Organic | 0.309 | 0.0089 | — | — | — | — | — |
| Organic | 0.309 | 0.0047 | Arachnida | Mesostigmata | Veigaiidae | Veigaia | Veigaia_nemorensis |
| Organic | 0.309 | 0.0061 | Collembola | Entomobryomorpha | — | — | — |
| Organic | 0.277 | 0.0282 | Arachnida | Trombidiformes | Tydeidae | — | — |
| Organic | 0.275 | 0.025 | Collembola | Entomobryomorpha | Isotomidae | — | — |
| Organic | 0.267 | 0.0276 | Arachnida | Sarcoptiformes | Phthiracaridae | Steganacarus | Steganacarus_applicatus |
| Organic | 0.267 | 0.0251 | Collembola | Entomobryomorpha | Isotomidae | Folsomia | Folsomia_quadrioculata |
| Organic | 0.267 | 0.0235 | — | — | — | — | — |
| Organic | 0.267 | 0.0324 | — | — | — | — | — |
| Organic | 0.267 | 0.0242 | Arachnida | Sarcoptiformes | Suctobelbidae | Suctobelbella | Suctobelbella_arcana |
| Organic | 0.267 | 0.0324 | — | — | — | — | — |
| Organic | 0.267 | 0.0276 | Insecta | — | — | — | — |
| Organic | 0.267 | 0.0276 | — | — | — | — | — |
| Organic | 0.267 | 0.0231 | Arachnida | Sarcoptiformes | Euphthiracaridae | Euphthiracarus | Euphthiracarus_monodactylus |
| Organic | 0.267 | 0.0243 | Arachnida | Sarcoptiformes | Phthiracaridae | Phthiracarus | Phthiracarus_clavatus |
| Organic | 0.267 | 0.0319 | — | — | — | — | — |
| Organic | 0.267 | 0.0208 | Arachnida | Sarcoptiformes | Phthiracaridae | Steganacarus | Steganacarus_spinosus |

### Previously clear-cut forests (CC)

| Association | Indicator value | Adjusted p | Class | Order | Family | Genus | Species |
| --- | --- | --- | --- | --- | --- | --- | --- |
| DW1 | 0.613 | 0.004 | Insecta | Coleoptera | Curculionidae | Pityogenes | Pityogenes_chalcographus |
| DW1 | 0.55 | 0.0173 | Insecta | Coleoptera | Curculionidae | Pityogenes | Pityogenes_chalcographus |
| DW1 | 0.47 | 0.0283 | — | — | — | — | — |
| DW1 | 0.449 | 0.0278 | — | — | — | — | — |
| DW1+DW2 | 0.624 | 0.0276 | Insecta | — | — | — | — |
| DW3 | 0.566 | 0.0276 | Insecta | — | — | — | — |
| DW3+DW4/5 | 0.674 | 0.004 | Arachnida | Sarcoptiformes | Carabodidae | Carabodes | — |
| DW3+DW4/5 | 0.485 | 0.0265 | Insecta | — | — | — | — |
| DW3+Organic | 0.461 | 0.029 | — | — | — | — | — |
| DW4/5 | 0.488 | 0.0417 | — | — | — | — | — |
| Organic | 0.707 | 0.004 | Arachnida | Sarcoptiformes | Nanhermanniidae | Nanhermannia | Nanhermannia_sellnicki |
| Organic | 0.671 | 0.004 | — | — | — | — | — |
| Organic | 0.661 | 0.004 | — | — | — | — | — |
| Organic | 0.652 | 0.004 | Collembola | Entomobryomorpha | Isotomidae | Isotomiella | Isotomiella_minor |
| Organic | 0.548 | 0.004 | Insecta | Coleoptera | Elateridae | Athous | Athous_subfuscus |
| Organic | 0.5 | 0.004 | Collembola | Poduromorpha | Neanuridae | Micranurida | Micranurida_forsslundi |
| Organic | 0.488 | 0.0064 | Collembola | Poduromorpha | Onychiuridae | Protaphorura | Protaphorura_pseudovanderdrifti |
| Organic | 0.468 | 0.0276 | Arachnida | Sarcoptiformes | Tectocepheidae | Tectocepheus | Tectocepheus_velatus |
| Organic | 0.447 | 0.0064 | Arachnida | Sarcoptiformes | Ceratozetidae | Ceratozetella | Ceratozetella_thienemanni |
| Organic | 0.428 | 0.029 | Insecta | — | — | — | — |
| Organic | 0.4 | 0.038 | — | — | — | — | — |
| Organic | 0.387 | 0.0295 | Insecta | — | — | — | — |
| Organic | 0.387 | 0.0295 | Insecta | Diptera | Sciaridae | — | — |
| Organic | 0.387 | 0.0295 | Collembola | Entomobryomorpha | Isotomidae | Folsomia | Folsomia_sensibilis |
| Organic | 0.387 | 0.0283 | Collembola | Entomobryomorpha | Isotomidae | Isotomiella | Isotomiella_minor |
| Organic | 0.387 | 0.029 | Collembola | Poduromorpha | Neanuridae | Anurida | Anurida_granulata |
| Organic | 0.387 | 0.0329 | Collembola | Entomobryomorpha | Isotomidae | Parisotoma | Parisotoma_notabilis |
| Organic | 0.387 | 0.0296 | — | — | — | — | — |
| Organic | 0.387 | 0.0295 | — | — | — | — | — |
| Organic | 0.387 | 0.0295 | Arachnida | Trombidiformes | Tydeidae | — | — |
| Organic | 0.387 | 0.029 | Arachnida | Sarcoptiformes | Oppiidae | Oppiella | Oppiella_nova |
| Organic | 0.387 | 0.029 | Insecta | — | — | — | — |

### Near-natural forests (NN)

| Association | Indicator value | Adjusted p | Class | Order | Family | Genus | Species |
| --- | --- | --- | --- | --- | --- | --- | --- |
| DW1 | 0.417 | 0.0322 | Insecta | Coleoptera | Curculionidae | Pityogenes | Pityogenes_chalcographus |
| DW1+DW2 | 0.579 | 0.0381 | Insecta | — | — | — | — |
| DW1+DW2 | 0.519 | 0.0468 | Insecta | — | — | — | — |
| DW1+DW2+DW3 | 0.798 | 0.0027 | Insecta | — | — | — | — |
| DW3+DW4/5 | 0.641 | 0.0064 | Arachnida | Sarcoptiformes | Carabodidae | Carabodes | — |
| DW3+DW4/5 | 0.576 | 0.0084 | — | — | — | — | — |
| DW4/5 | 0.433 | 0.0173 | Arachnida | Sarcoptiformes | Oribotritiidae | Protoribotritia | — |
| DW4/5+Organic | 0.534 | 0.0301 | — | — | — | — | — |
| DW4/5+Organic | 0.504 | 0.0227 | Collembola | Poduromorpha | Hypogastruridae | Willemia | Willemia_anophthalma |
| DW4/5+Organic | 0.463 | 0.0381 | Collembola | Neelipleona | Neelidae | Megalothorax | — |
| DW4/5+Organic | 0.459 | 0.03 | Collembola | Poduromorpha | Onychiuridae | Protaphorura | Protaphorura_pseudovanderdrifti |
| DW4/5+Organic | 0.459 | 0.0416 | Insecta | — | — | — | — |
| Organic | 0.916 | 0.0027 | Collembola | Entomobryomorpha | Isotomidae | Isotomiella | Isotomiella_minor |
| Organic | 0.798 | 0.0027 | — | — | — | — | — |
| Organic | 0.707 | 0.0027 | Arachnida | Sarcoptiformes | Nanhermanniidae | Nanhermannia | Nanhermannia_sellnicki |
| Organic | 0.634 | 0.0027 | Insecta | — | — | — | — |
| Organic | 0.603 | 0.0027 | Collembola | Entomobryomorpha | Isotomidae | Folsomia | Folsomia_sensibilis |
| Organic | 0.603 | 0.0027 | — | — | — | — | — |
| Organic | 0.564 | 0.0027 | Insecta | Diptera | Sciaridae | — | — |
| Organic | 0.522 | 0.0027 | — | — | — | — | — |
| Organic | 0.522 | 0.0027 | Collembola | Entomobryomorpha | Isotomidae | Isotomiella | Isotomiella_minor |
| Organic | 0.522 | 0.0027 | Arachnida | Sarcoptiformes | Oppiidae | Oppiella | Oppiella_nova |
| Organic | 0.522 | 0.0027 | Arachnida | Sarcoptiformes | Ceratozetidae | Ceratozetella | Ceratozetella_thienemanni |
| Organic | 0.504 | 0.0121 | — | — | — | — | — |
| Organic | 0.498 | 0.0068 | — | — | — | — | — |
| Organic | 0.477 | 0.0064 | Arachnida | Sarcoptiformes | Nothridae | Nothrus | Nothrus_silvestris |
| Organic | 0.477 | 0.0068 | Collembola | Poduromorpha | Neanuridae | Friesea | Friesea_mirabilis |
| Organic | 0.477 | 0.0064 | — | — | — | — | — |
| Organic | 0.477 | 0.0084 | Collembola | Poduromorpha | Neanuridae | Anurida | Anurida_granulata |
| Organic | 0.477 | 0.0081 | — | — | — | — | — |
| Organic | 0.477 | 0.0068 | Arachnida | Trombidiformes | Tydeidae | — | — |
| Organic | 0.477 | 0.0068 | Insecta | Diptera | Bibionidae | Bibio | — |
| Organic | 0.426 | 0.0167 | Insecta | — | — | — | — |
| Organic | 0.426 | 0.0084 | — | — | — | — | — |
| Organic | 0.426 | 0.023 | — | — | — | — | — |
| Organic | 0.424 | 0.0332 | Arachnida | Sarcoptiformes | Tectocepheidae | Tectocepheus | Tectocepheus_velatus |
| Organic | 0.419 | 0.035 | Arachnida | Trombidiformes | Eupodidae | — | — |
| Organic | 0.413 | 0.0312 | Arachnida | Sarcoptiformes | Crotoniidae | Platynothrus | Platynothrus_peltifer |
| Organic | 0.369 | 0.0402 | Collembola | Poduromorpha | Hypogastruridae | Willemia | Willemia_anophthalma |
| Organic | 0.369 | 0.0368 | Insecta | — | — | — | — |
| Organic | 0.369 | 0.0376 | — | — | — | — | — |
| Organic | 0.369 | 0.0388 | Collembola | Entomobryomorpha | — | — | — |
